# The landscape of m1A modification and its posttranscriptional regulatory functions in primary neurons

**DOI:** 10.1101/2023.01.25.525483

**Authors:** Chi Zhang, Xianfu Yi, Mengfan Hou, Qingyang Li, Xueying Li, Lu Lu, Enling Qi, Mingxin Wu, Lin Qi, Huan Jian, Zhangyang Qi, Yigang Lv, Xiaohong Kong, Mingjun Bi, Shiqing Feng, Hengxing Zhou

## Abstract

Cerebral ischaemia‒reperfusion injury, during which neurons undergo oxygen-glucose deprivation/reoxygenation (OGD/R), is a notable pathological process in many neurological diseases. N1-methyladenosine (m1A) is an RNA modification that can affect gene expression and RNA stability. The m1A landscape and potential functions of m1A modification in neurons remain poorly understood. We explored RNA (mRNA, lncRNA, and circRNA) m1A modification in normal and OGD/R-treated neurons and the effect of m1A on diverse RNAs. We investigated the m1A landscape in primary neurons, identified m1A-modified RNAs, and found that OGD/R increased the number of m1A RNAs. m1A modification might also affect the regulatory mechanisms of noncoding RNAs, e.g., lncRNA–RBP interactions and circRNA translation. We showed that m1A modification mediates the circRNA/lncRNA‒miRNA–mRNA ceRNA mechanism and that 3’UTR methylation of mRNAs can hinder miRNA–mRNA binding. Three methylation patterns were identified, and genes with different patterns had intrinsic mechanisms with potential m1A-regulatory specificity. Systematic analysis of the m1A landscape in normal and OGD/R neurons lays a critical foundation for understanding RNA methylation and provides new perspectives and a theoretical basis for treating and developing drugs for OGD/R pathology-related diseases.

## INTRODUCTION

RNA modifications were first identified more than 50 years ago *(**Dunn, 1961**)*. With the development of sequencing technologies, our understanding of the features (location, function, and regulation) of RNA modifications has greatly improved. RNA modifications include N6-methyladenosine (m6A), N1-methyladenosine (m1A), 5-hydroxymethylcytosine (hm5C), 5-methylcytosine (m5C), pseudouridine (Ψ), and inosine *(**Yoon et al., 2017**)*. Several studies have indicated that in addition to m6A, m1A is an abundant epitranscriptomic modification and regulates multiple biological processes ranging from local structural stability *(**Oerum et al., 2017**)* to RNA–protein interactions *(**Zhao et al., 2019**)*, apoptosis and cell proliferation *(**Li et al., 2016; Chen et al., 2019c**)*.

m1A was first differentiated from other RNA modifications by Dunn in 1961 *(**Dunn, 1961**)*. m1A modification constitutes the addition of a methyl group to the N1 position of adenosine via m1A regulators. Similar to m6A RNA methylation, three kinds of regulators mediate m1A status: ‘writers’ (TRMT10C, TRMT61B, and TRMT6/61A) *(**Chujo and Suzuki, 2012; Engel and Chen, 2018**)*, ‘readers’ (YTHDF1, YTHDF2, YTHDF3, and YTHDC1) *(**Dai et al., 2018; Seo and Kleiner, 2020**)*, and ‘erasers’ (ALKBH1 and ALKBH3) *(**Liu et al., 2016; Safra et al., 2017**)*. Writers are methyltransferases that manipulate the level of m1A to interfere with translation. Human mitochondrial tRNAs (mt-tRNAs) contain m1A at positions 9 and 58 *(**Guelorget et al., 2010**)*. TRMT61B and TRMT6/61A catalyse m1A modification at position 58 in mt-RNA in human cells, while TRMT10C catalyses it at position 9 *(**Ozanick et al., 2005; Chujo and Suzuki, 2012**)*. Readers recognize m1A and mediate the translation and degradation of downstream RNAs. Proteins containing YTH domains directly bind to m1A-modified positions in RNA transcripts *(**Dai et al., 2018**)*. Erasers are demethylases that catalyse the removal of m1A from single-stranded DNA and RNA *(**Liu et al., 2016**)*. Two AlkB family proteins, ALKBH3 and ALKBH1, have been found to remove m1A *(**Ueda et al., 2017**)*.

Cerebral ischaemia‒reperfusion injury (IRI) is the main pathological process in many brain diseases, such as traumatic brain injury (TBI) *(**Zhao et al., 2018**)* and acute ischaemic stroke *(**Wiberg et al., 2016**)*. Ischaemia and blood flow reperfusion cause damage at ischaemic sites. Neuronal oxygen-glucose deprivation/reoxygenation (OGD/R), which results in dysregulation of material and energy metabolism, subsequently affects neuronal biological processes such as survival *(**Sasaki et al., 2011**)*, apoptosis *(**Song et al., 2019**)*, and autophagy *(**Wang et al., 2014**)*. Dysfunction of biological processes resulting from OGD/R leads to the occurrence and aggravation of diseases *(**Wang et al., 2014; Wiberg et al., 2016; Song et al., 2019**)*. Therefore, it is highly important to study the changes in neurons undergoing OGD/R. Recently, many studies have shown that various mRNAs and noncoding RNAs (long noncoding RNAs [lncRNAs], circular RNAs [circRNAs], and microRNAs [miRNAs]) regulate the exacerbation or amelioration of pathological processes caused by OGD/R *(**Wang et al., 2017; Chen et al., 2020, 2021; Yang et al., 2021**)* through various mechanisms. For example, the lncRNA U90926 directly binds to malate dehydrogenase 2 (MDH2), which could aggravate ischaemic brain injury by facilitating neutrophil infiltration *(**Chen et al., 2021**)*. circUCK2 functions as a sponge to inhibit miR-125b-5p activity, resulting in an increase in growth differentiation factor 11 (GDF11) expression and subsequent amelioration of neuronal injury *(**Yang et al., 2021**)*. Some studies have shown that chemical modifications of these RNAs can regulate the original mechanism *(**Patil et al., 2016; Yang et al., 2018; Chen et al., 2019a; Wen et al., 2020; Xu et al., 2020; Liu et al., 2021b**)*. However, most studies have focused on m6A modification, and little is known about the features and functions of m1A modification of various RNAs in normal neurons and OGD/R-treated neurons.

In this study, we identified the characteristics of m1A modification of various RNAs (mRNA, lncRNA, and circRNA) in neurons and explored the potential effects of these modifications on different RNA functions. We first identified m1A-modified peaks (m1A peaks) in normal neurons and OGD/R-treated neurons at different times and thus discovered a GA-rich motif. The number of RNAs with m1A modification, the chromosome distribution, and the changes in m1A modification on various RNAs before and after OGD/R treatment were analysed. The number of m1A RNAs was increased after OGD/R treatment, and most of these differentially expressed m1A-modified RNAs were involved in biological processes and signalling pathways related to the regulation of cellular homeostasis and synapses. In addition, the results indicated that m1A modification mediates the complex posttranscriptional regulation network and that the regulation of m1A modification in key nodes of RNA interaction networks may cause widespread changes in downstream signalling. We also proposed three patterns of m1A methylation, with genes fitting these methylation patterns participating in different biological functions and signalling pathways. Overall, we analysed the characteristics of m1A modification of multiple RNAs in normal neurons and OGD/R-treated neurons from multiple perspectives. The landscape and other findings provide evidence useful for the exploration of epitranscriptomic mechanisms and the development of targeted drugs in OGD/R-related pathological processes.

## RESULTS

### The common features of m1A modification in primary neurons

The distribution of m1A modifications in the transcriptome is uncharacterized in the nervous system, especially in important component neurons. We performed MeRIP-seq to clarify the m1A transcriptomic landscape in primary neurons and OGD/R-treated neurons (Figure 1A). A total of 48260, 13588, and 16397 m1A peaks were identified in mRNAs, lncRNAs, and circRNAs, respectively (Figure 1B). For example, regarding the m1A peaks in mRNAs, 29066, 37893, and 41334 m1A peaks were identified in the control, OGD/R 1.5 h, and OGD/R 3 h samples, respectively (Figure supplement 1A). We examined the genomic distribution of those m1A peaks and found that in the control group, 54.2% were located in CDSs and 25.1% were located in the 5’UTRs in mRNAs, with 88.4% located in CDSs and 8.9% in 5’UTRs in circRNAs (Figure 1C). A comparable pattern was found in the OGD/R 1.5 h and 3 h groups (Figure supplement 1B). The genomic distribution pattern was slightly different from previous findings *(**Dominissini et al., 2016**)*. However, more m1A peaks were in the CDS in circRNAs than in mRNAs (88.4% vs. 54.2% in the control group), and a clear decreasing trend in near-5’UTRs was observed (Figure 1D and Figure supplement 1C). The m1A genomic distribution in mRNAs and circRNAs differed significantly between normal and OGD/R-treated neurons (*chi*-squared test*, p < 0.05*, Figure supplement 1B). The above results indicate the presence of abundant m1A modifications in neurons and show that different OGD/R treatments may affect these modifications.

**Figure 1.**
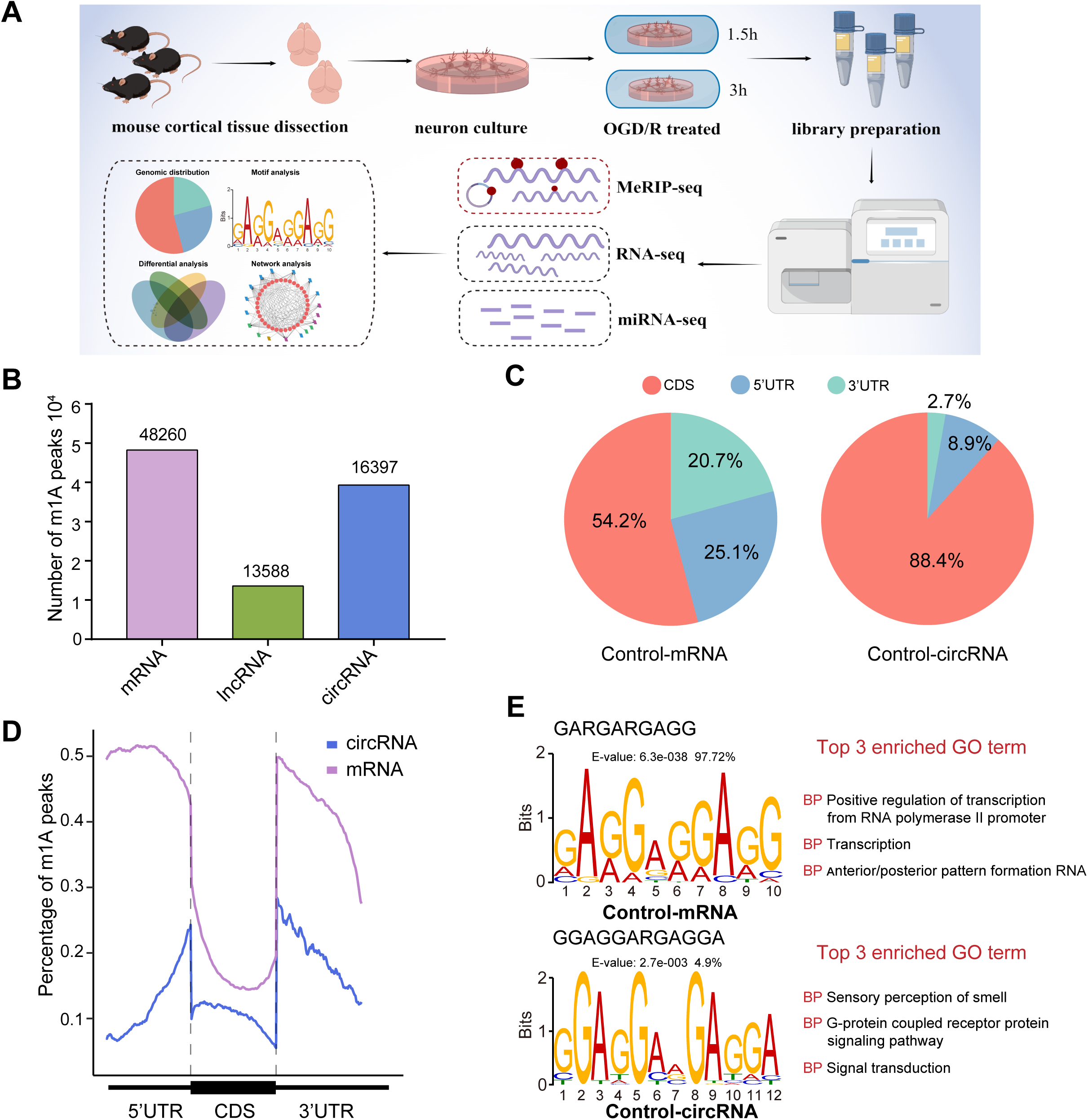
The common m1A modification features in primary neurons. A. Schematic of the experimental design and data analysis workflow (drawn by Figdraw). B. m1A peaks identified in different RNAs (mRNA, lncRNA, and circRNA). C. The genomic distribution of m1A peaks in mRNAs and circRNAs. D. The genomic distribution pattern of m1A peaks in mRNAs and circRNAs on a metagene. E. Potential motifs for m1A peaks in both m1A mRNAs and m1A circRNAs in the control group. The E-value of a motif is based on its log likelihood ratio, width, sites, and background letter frequencies. The percentage represents the proportion of a motif in all identified motifs. **Figure supplement 1. The common m1A modification features in primary neurons**

Some studies have shown that some m1A motifs, such as GUUCNANNC motifs *(**Safra et al., 2017**)*, GUUCRA motifs *(**Li et al., 2017**)*, and GA-rich consensus sequences *(**Li et al., 2016**)*, exist in different cells. However, there are no accepted highly conserved regions for m1A modification. We used unbiased motif detection to reveal the potential motifs for m1A peaks with MEME Suite *(**Bailey et al., 2015**)* (v5.4.1). Various motifs that were also GA-rich were identified in mRNAs and circRNAs (Figure 1E and supplement 1D). Indeed, more than 90% (4510/4646 in the control group) of motifs in mRNA were GA-rich, and approximately 10% (579/4391 in the control group) of motifs in circRNA were GA-rich. This pattern indicated that GA-rich m1Amotifs are more prevalent in mRNA and that various mechanisms may control m1A modification in circRNA. By comparing these motifs with the known motifs in the JASPAR database *(**Castro-Mondragon et al., 2022**)* (v2022), we found that the GA-rich motifs are highly similar to those in MA0528.1 from ZNF263 (Figure 1E and supplement 1D), and the functions of these motifs were mainly focused on RNA polymerase II promoter regulation. Interestingly, although the motifs in circRNAs in the OGD/R 3 h group were also GA-rich, their main function was to regulate axon guidance and the G-protein coupled receptor protein signalling pathway. These results suggest that GA-rich m1A motifs are present in neurons, although they vary across RNAs with different functions.

### OGD/R increases the number of m1A mRNAs and affects neuron fate

m6A modifications can functionally alter the expression of mRNAs, pre-mRNAs, miRNAs, and noncoding RNAs, such as rRNAs and tRNAs *(**Chua et al., 2020**)*. Although m1A is another abundant RNA modification, the characteristics of this modification on mRNAs and noncoding RNAs in neurons remain unclear. We identified m1A modifications on different kinds of RNAs (mRNAs, lncRNAs, and circRNAs) in normal primary neurons and OGD/R-treated neurons to determine whether there are characteristic modifications of different RNAs after OGD/R treatment. First, we determined the numbers of m1A mRNAs identified by four different methods: the conventional, treatment, mismatch, and trough methods (see the Materials and methods section for details). The most m1A mRNAs were identified by the conventional method, while the mismatch method identified the least (Figure supplement 2A). We also identified the unique and common m1A mRNAs identified in different OGD/R-treated neurons by 4 different methods (Figure supplement 2B). The percentages of common (5.82%-76.44%) and unique mRNAs (8.48%-70.1% in the control group, 11.56%-74.17% in the OGD/R 1.5 h group, and 11.59%-76.03% in the OGD/R 3 h group) identified by the different methods varied greatly. To ensure the accuracy of subsequent analyses as much as possible, we selected the m1A mRNAs identified by at least two of the abovementioned methods for subsequent analyses (Figure supplement 2C), resulting in 4047, 5756, and 4948 (Source data1) m1A mRNAs for the control, OGD/R 1.5 h, and OGD/R 3 h groups, respectively (Figure 2A).

**Figure 2.**
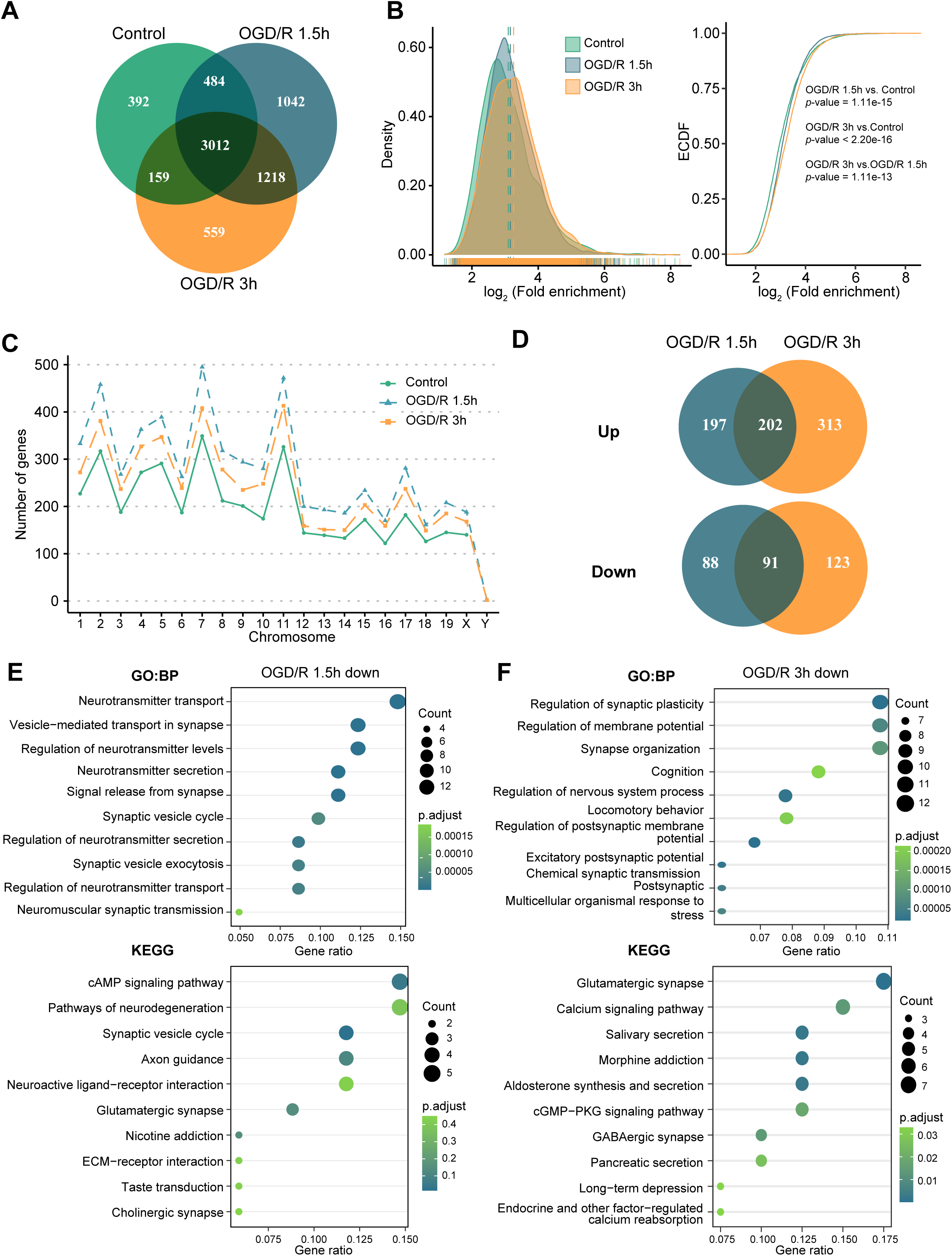
OGD/R increases the number of m1A mRNAs and affects neuron fate. A. Shared and unique m1A mRNAs in the control and different OGD/R groups. B. The density distribution (left) and cumulative distribution function curves (right) to show the m1A modification level among different groups. The Kolmogorov‒Smirnov test was used to test the significance. C. The number of m1A mRNAs on each chromosome in different groups. D. Venn diagram showing differentially methylated (up and down) m1A mRNAs in the OGD/R 1.5 h and OGD/R 3 h groups. E-F. GO and KEGG analyses of differentially methylated (down) m1A mRNAs in the OGD/R 1.5 h (E) and OGD/R 3 h (F) groups. **Figure supplement 2. OGD/R increases the number of m1A mRNAs and affects neuron fate Source data 1** **Source data 2**

Then, we used conventional density distribution (Figure 2B, left) and cumulative distribution function curves (Figure 2B, right) to compare the m1A level among the groups. After OGD/R treatment, the m1A level in neurons was higher than that in the control group (OGD/R 1.5 h vs. control: *p* < 1.1e-15, OGD/R 3 h vs. control: *p* < 2.2e-16, Kolmogorov‒Smirnov test). Since the modification density differed by treatment, we sought to determine whether the amount of m1A modification on each chromosome differs before and after treatment. The numbers of m1A mRNAs on each chromosome were increased after OGD/R treatment (Figure 2C).

Finally, differential methylation analysis was conducted between OGD/R-treated neurons and normal neurons (Source data2). During the different OGD/R treatments, the mRNAs with large changes in methylation levels were not totally consistent between the two groups, possibly implying that distinct m1A modification patterns exist in the different OGD/R-treated groups (Figure 2D). To further clarify the functions of the differentially m1A methylated mRNAs, GO and KEGG enrichment analyses were performed. The m1A mRNAs with decreased methylation in the OGD/R 1.5 h group regulated synapses, the release of neurotransmitters, the cAMP signalling pathway, and axon guidance (Figure 2E). In contrast, the m1A mRNAs with decreased methylation in the OGD/R 3 h group regulated synaptic plasticity, the membrane potential, glutamatergic synapses, and the calcium signalling pathway (Figure 2F). The m1A mRNAs with increased methylation in the OGD/R 1.5 h group influenced morphogenesis, cell fate commitment, chemical carcinogenesis and apoptosis, while the m1A mRNAs with increased methylation in the OGD/R 3 h group affected transmembrane receptors, cell substrate adhesion, focal adhesion, and regulation of the actin cytoskeleton (Figure supplement 2D and 2E). The above results indicate that under different OGD/R treatments, the increases and decreases in m1A in neurons affect different biological functions. With prolonged OGD/R time, differential m1A modification regulates the fate of neurons.

### OGD/R increases the number of m1A lncRNAs and affects RNA processing

m6A modification has been reported to be present on lncRNAs and to affect their biological functions *(**Patil et al., 2016; Yang et al., 2018; Wen et al., 2020**)*. However, the m1A modification of lncRNAs and its possible role in neurons remain uncharacterized. We identified lncRNAs with m1A modification in normal neurons and OGD/R-treated neurons to explore the potential functions of m1A lncRNAs. First, we applied the 4 abovementioned methods to identify m1A lncRNAs in normal and OGD/R-treated neurons. The conventional method identified the most m1A lncRNAs (Figure supplement 3A). We then identified the unique and common m1A lncRNAs in different OGD/R-treated neurons by the 4 different methods (Figure supplement 3B). Fewer m1A lncRNAs than m1A mRNAs were identified by the 4 methods (Figure supplement 3C). Similarly, the m1A lncRNAs identified by at least two methods were selected for further analysis (control: 1078, OGD/R 1.5 h: 1661, OGD/R 3 h: 1294) (Figure 3A, Figure supplement 3D and Source data3).

**Figure 3.**
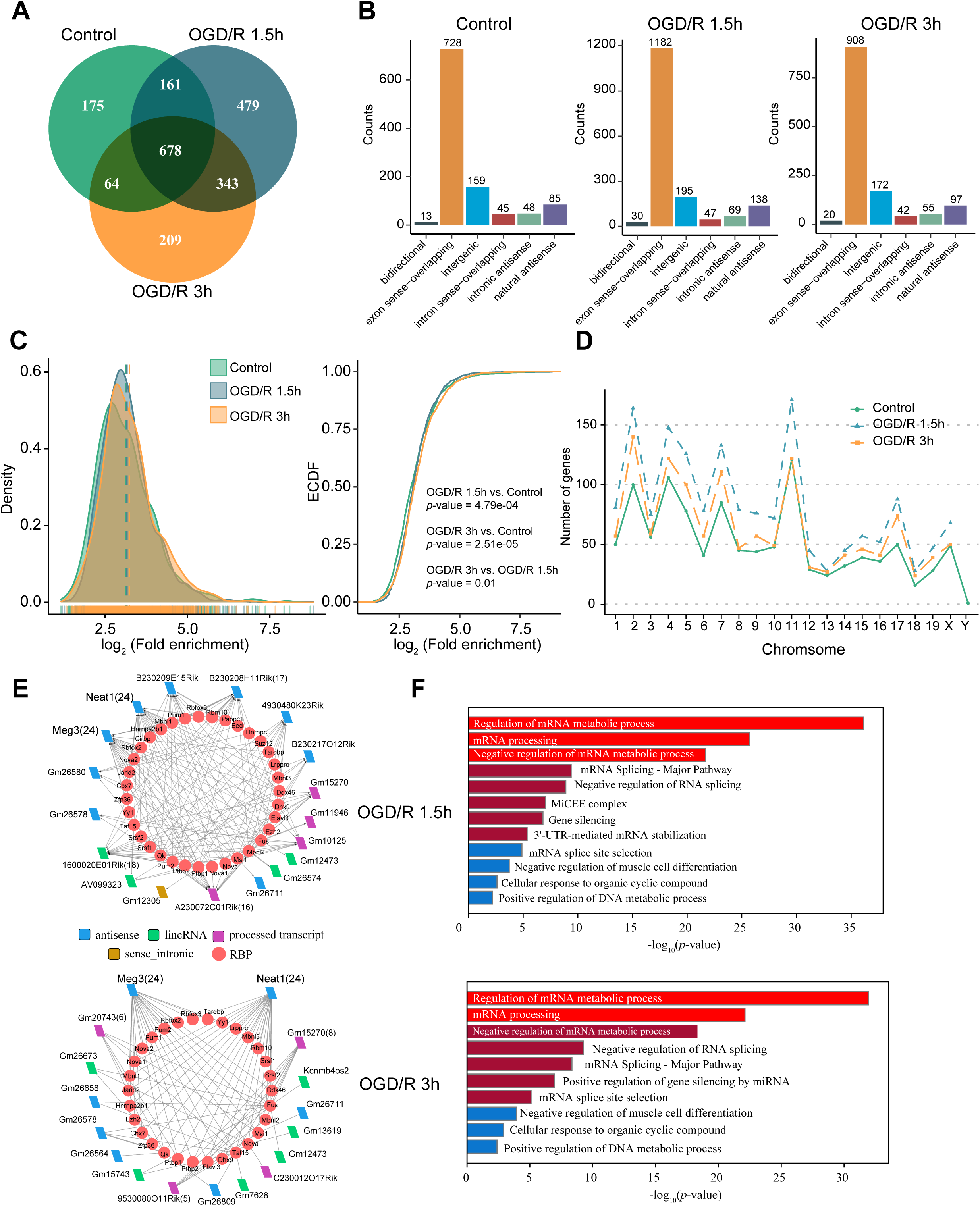
OGD/R increases the number of m1A lncRNAs and affects RNA processing. A. Shared and unique m1A lncRNAs in the control and different OGD/R groups. B. The genomic resources of m1A lncRNAs in the control and OGD/R-treated groups. C. The density distribution (left) and cumulative distribution function curves (right) show the m1A modification level among the different groups. The Kolmogorov‒Smirnov test was used to test the significance. D. The number of m1A lncRNAs on each chromosome in different groups. E-F. The interaction networks between lncRNAs and RBPs in different OGD/R groups (E) and functional enrichment analysis of those core RBPs (F). **Figure supplement 3. OGD/R increases the number of m1A lncRNAs and affects RNA processing** **Source data 3** **Source data 4**

We next analysed the genomic sources of m1A lncRNAs in the control and OGD/R-treated groups. The two main genomic sources of m1A lncRNAs were exon sense-overlapping regions and intergenic regions (Figure 3B). This distribution did not differ significantly between the control and OGD/R groups (*chi*-squared test, *p* = 0.303). Similar to the mRNA modification characteristics (Figure 2B), the differences in the lncRNA m1A modification level between the control and OGD/R-treated groups were statistically significant (Figure 3C, OGD/R 1.5 h vs. control: *p* < 0.001, OGD/R 3 h vs. control: *p* < 2.508e-5; Kolmogorov‒Smirnov test). We also counted the m1A lncRNAs on each chromosome in normal and OGD/R-treated neurons and found that the number of m1A lncRNAs on each chromosome was increased after OGD/R treatment (Figure 3D).

Interaction with RNA binding proteins (RBPs) is an important regulatory mechanism of lncRNAs *(**Yang et al., 2020a; Søndergaard et al., 2022**)*. We thus sought to determine whether lncRNAs with m1A modifications play vital roles in biological processes, such as RNA processing. Considering the differentially m1A-methylated lncRNAs in the OGD/R groups (Figure supplement 3E) (Source data4), RBPs binding to these lncRNAs with high reliability were predicted in the ENCORI database, and lncRNA–RBP interaction networks were then constructed (Figure 3E). In the OGD/R 1.5 h and OGD/R 3 h lncRNA–RBP networks, Meg3 and Neat1, which have been reported to play an important role in nervous system-related diseases, were predicted to bind to various RBPs *(**Zhong et al., 2017; Sanli et al., 2018; Cui et al., 2019; Liang et al., 2020**)*. Functional enrichment analysis showed that these RBPs participated in the processes of mRNA metabolism, mRNA processing, and mRNA splicing (Figure 3F). These results suggest that the ability of nerve-related lncRNAs to regulate RNA metabolism and then affect the pathophysiological processes in neurological diseases may depend on their m1A modification.

### OGD/R increases the number of m1A modification sites in circRNAs and regulates translation functions

As RNA molecules regulating diverse pathophysiological processes, circRNAs are specifically enriched in the nervous system *(**Gokool et al., 2020; Mehta et al., 2020**)*. Some studies have demonstrated that m6A modification can affect circRNA biogenesis, immunogenicity, translation, etc. *(**Chen et al., 2019b; Tang et al., 2020; Xu et al., 2020**)*. However, the features of m1A modification of circRNAs in the nervous system remain unknown. We investigated m1A modifications in neuronal circRNAs using various approaches. The four abovementioned methods identified different numbers of m1A circRNAs in each group, with the most m1A circRNAs identified by the conventional method (Figure supplement 4A). We then examined the common and unique m1A circRNAs in the normal and OGD/R-treated groups identified by the 4 methods. The fewest associated m1A circRNAs were identified by the mismatch method (Figure supplement 4A and 4B). Fewer m1A circRNAs than m1A mRNAs were identified by all 4 methods (Figure supplement 4C). However, m1A circRNAs were more abundant than m1A lncRNAs. To further investigate the role of m1A circRNAs in each group, we selected the m1A circRNAs identified by at least two methods for downstream analysis (Figure 4A, Figure supplement 4D and Source data5).

**Figure 4.**
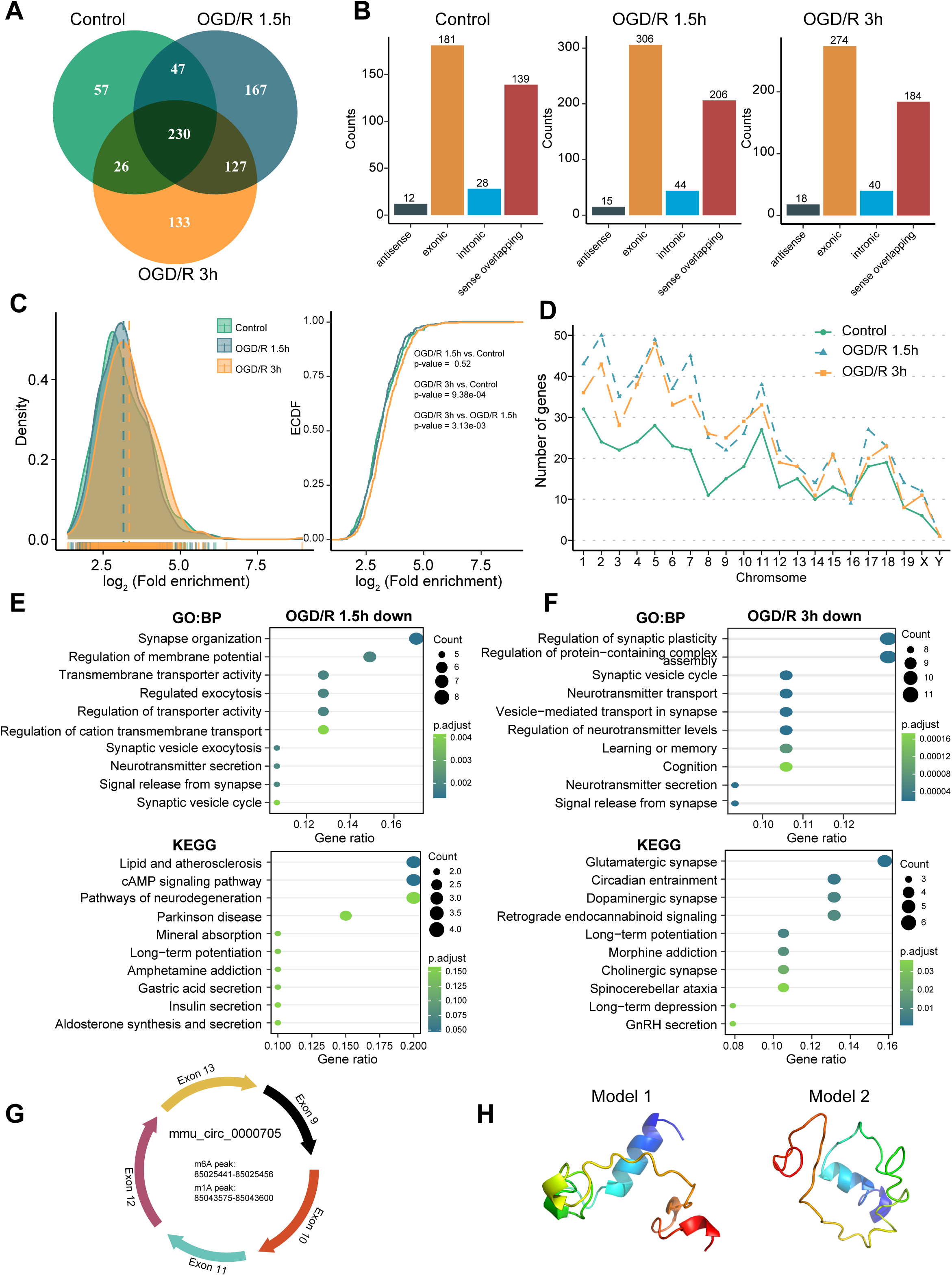
OGD/R increases the m1A modification sites on circRNA and regulates translation functions. A. Shared and unique m1A circRNAs in the control and different OGD/R groups. B. The genomic resources of m1A circRNAs in the control and OGD/R-treated groups. C. The density distribution (left) and cumulative distribution function curves (right) show the m1A modification level among the different groups. The Kolmogorov‒Smirnov test was used to test the significance. D. The number of m1A circRNAs on each chromosome in different groups. E-F. GO and KEGG analysis of differentially methylated (down) m1A circRNAs in OGD/R 1.5 h (E) and OGD/R 3 h (F). G. An example of a circRNA (mmu_circ_0000705) with translation ability that also contains an m1A and m6A modification site. H. Predicted polypeptide structure of mmu_circ_0000705. **Figure supplement 4. OGD/R increases the m1A modification sites on circRNA and regulates translation functions** **Source data 5** **Source data 6**

The genomic sources of those m1A circRNAs were counted in each group, and exonic regions and sense-overlapping regions were found to be the main sources (*chi*-squared test, *p* = 0.9379) (Figure 4B). Next, we generated density distribution and cumulative distribution function curves to explore the m1A modification degree in the three groups. The m1A modification level was increased after OGD/R treatment, and the difference between the OGD/R 3 h and control groups was statistically significant (OGD/R 3 h vs. control *p* < 0.0009, Kolmogorov‒Smirnov test). However, the m1A level did not differ significantly between the OGD/R 1.5 h and control groups (Figure 4C). m1A circRNAs on each chromosome were also analysed, and the numbers of m1A circRNAs were increased after the different OGD/R treatments (Figure 4D).

To clarify the biological functions of the differentially m1A-methylated circRNAs in the OGD/R 1.5 h and OGD/R 3 h groups (Source data6), we obtained the source genes of these differentially methylated circRNAs and performed GO and KEGG functional enrichment analyses. In the OGD/R 1.5 h group, the differentially methylated circRNAs with decreased m1A levels were enriched in synapse organization, regulation of membrane potential, the cAMP signalling pathway and the neurodegeneration pathway, while the circRNAs with increased m1A levels were enriched in morphogenesis, GTPase activity, axon guidance and the MAPK signalling pathway (Figure 4E and Figure supplement 4E). In the OGD/R 3 h group, the differentially methylated circRNAs with decreased m1A levels were enriched in synaptic plasticity, neurotransmitter transport, glutamatergic synapse and dopaminergic synapse, while the circRNAs with increased m1A levels were enriched in supramolecular fibre organization, cellular component disassembly, adherens junction and endocytosis (Figure 4F and Figure supplement 4F). These results indicate that the differentially modified circRNAs regulate different biological functions in different OGD/R processes. The m1A circRNAs with decreased m1A levels were related mainly to synapses and the release of neurotransmitters, while the m1A circRNAs with increased m1A levels played regulatory roles in many biological processes.

Currently, research on the translation ability and products of circRNAs is gradually increasing. We asked whether the m1A circRNAs could be translated. By using the riboCIRC database *(**Li et al., 2021**)* (http://www.ribocirc.com/index.html), we identified some circRNAs (237/811) with translation ability among the identified differentially m1A-modified circRNAs. In addition, m6A modifications annotated in the database were present on some m1A circRNAs. The polypeptide structures were predicted to show the possible translation products. Then, mmu_circ_0000705 (encoded by App) and mmu_circ_0002207 (encoded by Foxo3) were selected to show the abovementioned features (Figure 4G and H, Figure supplement 4G and 4H). App can regulate the stability of synapses that bind to diverse proteins and regulates the occurrence and development of nervous system diseases *(**Lee et al., 2020; Eysert et al., 2021**)*. mmu_circ_0000705 is composed of 5 exons. In this circRNA, m6A modification occurs in the 85025441-85025456 region, while m1A modification occurs in the 85043575-85043600 region. Foxo3 also regulates multiple functions of the nervous system *(**Deng et al., 2018; Du et al., 2021**)*. mmu_circ_0002207 is composed of 1 exon, with m6A modification occurring in the 42196615-42196629 region and m1A modification occurring in the 42196961-42197760 region. These results suggest that m1A modification may play a regulatory role in the translation of circRNAs into peptides. However, the specific mechanism needs further exploration.

### m1A modification affects the ceRNA mechanism of differentially methylated lncRNAs and circRNAs

The ceRNA mechanism is the main posttranscriptional regulatory mechanism of circRNAs and lncRNAs. Studies have shown that the ceRNA mechanism plays an important role in nervous system diseases *(**Huang et al., 2020; Moreno-Garcia et al., 2020**)*. However, the function and physiological effect of the m1A modifications in these noncoding RNAs remain unclear. We identified the differentially expressed miRNAs in the OGD/R-treated groups; 81 differentially modified miRNAs were found in the OGD/R 1.5 h group, and 69 differentially modified miRNAs were found in the OGD/R 3 h group (Figure 5A and Figure supplement 5A). Differentially expressed mRNAs in the OGD/R-treated groups were also identified by RNA-seq. A total of 1579 and 3259 differentially expressed mRNAs were identified in the OGD/R 1.5 h and OGD/R 3 h groups, respectively (Figure 5B and Figure supplement 5B). GO and KEGG enrichment analyses were conducted to explore the biological functions of the differentially expressed mRNAs (Figure 5C). In the OGD/R 1.5 h group, the differentially expressed mRNAs were enriched in extracellular matrix organization, the PI3K−Akt signalling pathway, neuroactive ligand−receptor interaction, and the calcium and Hippo signalling pathways. In the OGD/R 3 h group (Figure supplement 5C), the differentially expressed mRNAs were enriched in synapse organization, axonogenesis, regulation of membrane potential, regulation of neurogenesis, etc. The functions of the differentially expressed genes were related mainly to the influence on cell structure in the OGD/R 1.5 h group but to synapse, axon and other neuron functions in the OGD/R 3 h group.

**Figure 5.**
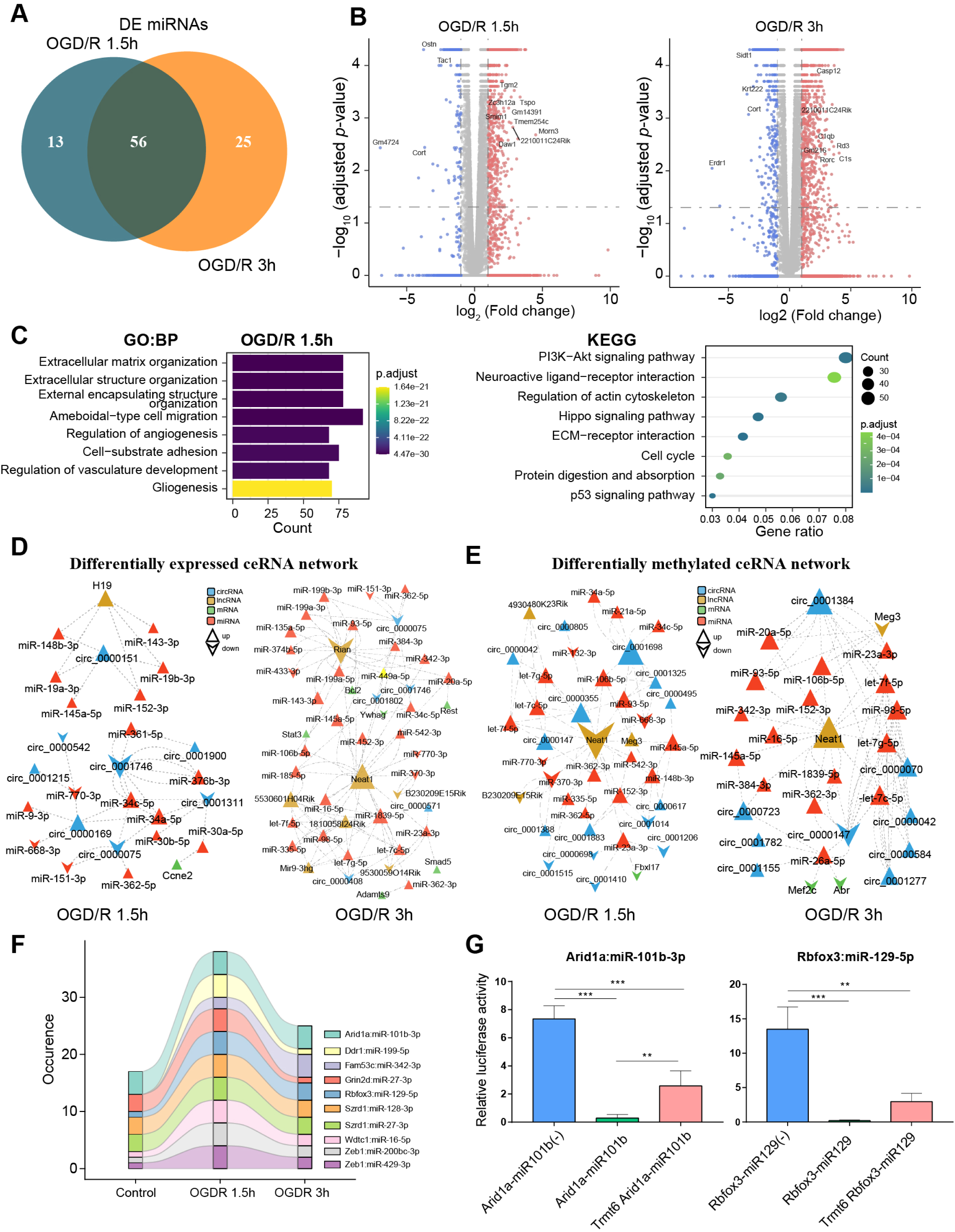
m1A modification affects the ceRNA mechanism of differentially methylated lncRNAs and circRNAs. A. Venn diagram showing differentially expressed miRNAs in different OGD/R-treated groups. B. Volcano plot showing differentially expressed mRNAs in different OGD/R-treated groups (OGD/R 1.5 h, left and OGD/R 3 h, right). C. GO analysis shows the biological functions of the differentially expressed mRNAs in the OGD/R 1.5 h group. D-E. Expression-(D) and methylation-specific (E) ceRNA regulatory networks in the OGD/R 1.5 h and OGD/R 3 h groups. F. Sankey diagram showing the dynamic changes in miRNA‒mRNA pairs in the control, OGD/R 1.5 h and OGD/R 3 h groups. G. A dual luciferase assay showed that m1A modification of Arid1a and Rbfox3 blocks the binding of the corresponding miRNAs. **Figure supplement 5. m1A modification affects the ceRNA mechanism of differentially methylated lncRNAs and circRNAs** **Source data 7** **Source data 8**

Based on the differentially expressed miRNAs (Source data7), we performed a screen to identify the differentially expressed and differentially methylated mRNAs, lncRNAs and circRNAs that bind to these miRNAs. Then, the identified miRNAs, mRNAs, lncRNAs and circRNAs were used to construct expression- and methylation-specific ceRNA regulatory networks. In the OGD/R 1.5 h expression ceRNA network, H19, a well-known lncRNA, was shown to regulate circRNAs by sponging miRNAs (Figure 5D, left and EV5D, top). In the OGD/R 3 h expression ceRNA network, Neat1 and Rian were shown to regulate various miRNAs and affect downstream mRNA and circRNA expression (Figure 5D, right and Figure supplement 5D, bottom). In the OGD/R 1.5 h methylation ceRNA network, Neat1, which plays critical regulatory roles in diverse neurological diseases, was shown to sponge many miRNAs and affect other lncRNA/circRNA–miRNA axes (Figure 5E, left and Figure supplement 5E). In the OGD/R 3 h methylation ceRNA network, Neat1 was also shown to play important roles (Figure 5E, right). Interestingly, the miRNAs sponged by Neat1 in the methylation ceRNA network were different from those in the expression ceRNA network. Neat1, present in both the expression and methylation ceRNA networks, sponged different miRNAs in the two networks, suggesting that m1A modification of lncRNAs impacts the ceRNA mechanism.

### m1A methylation of mRNA 3’UTRs hinders miRNA‒mRNA binding

Our above analysis indicated that m1A modification may affect the ceRNA mechanism of different RNAs. miRNA binding sites in mRNAs are located mainly in the 3’UTR. Therefore, we next explored the changes in miRNA‒mRNA pairs under different OGD/R treatments. By analysing the m1A-modified peak regions and the positions of miRNA binding seed sequences, we found that there was some miRNA‒mRNA pairs that changed dynamically during different OGD/R treatments (Figure 5F). Among these miRNA‒mRNA pairs, some mRNAs have been shown to be associated with neural tissue development (Rbfox3) *(**Kim et al., 2013**)*, neural progenitor differentiation (Arid1a) *(**Liu et al., 2021a**)*, NMDA receptor complexes (Grin2d) *(**XiangWei et al., 2019**)*, and ubiquitination of proteins (Wdtc1) *(**Groh et al., 2016**)*. Therefore, for these key miRNA‒mRNA pairs, we designed a dual luciferase assay to detect whether m1A modification of the 3’UTR affects the binding of miRNAs to mRNAs (Source data8). Arid1a encodes a member of the SWI/SNF family that belongs to the neural progenitor-specific chromatin remodelling complex (npBAF complex) and the neuron-specific chromatin remodelling complex (nBAF complex). We cotransfected Arid1a, miR101b and Trmt6 (an m1A methyltransferase) into HEK293T cells. Trmt6 alleviated the inhibitory effect of miRNA101b on Arid1a expression (one-way ANOVA, *p*<0.05). Rbfox3–miR129 is another miRNA‒mRNA pair affected by Trmt6. After cotransfection of Rbfox3–miR129 and Trmt6, Trmt6 alleviated the repression of target genes by the miRNA (albeit no significantly; one-way ANOVA) (Figure 5G). Regarding the other two genes (Grin2d and Wdtc1), after cotransfection with their corresponding miRNAs and Trmt6, Trmt6 did not affect the ability of the miRNAs to bind to their target genes (Figure supplement 5F). This result suggests that m1A methylation of the 3’UTR in some mRNAs does affect the binding of miRNAs. The underlying mechanisms appear complex and need further exploration.

### Three patterns of m1A modification regulation in neurons

We identified features of m1A modifications in mRNAs and noncoding RNAs and explored the potential functions of those RNAs. However, the m1A patterns remain unknown. We applied the NMF method to identify m1A modification patterns; *k* = 3 was the previous value at which the largest change occurred and was thus selected as the optimal *k* value (Figure supplement 6A). Three m1A modification patterns, termed Cluster 1, Cluster 2, and Cluster 3, were discovered (Figure 6A). The mixture coefficient matrix also showed that the three clusters had good discrimination (Figure 6B). We further explored whether the different m1A clusters had different biological functions. Cluster 1 was defined as the “metabolism-associated cluster” (MAC) since its members were enriched in signal transduction and biosynthetic processes (Figure 6C and Figure supplement 6B) and metabolism-associated pathways, such as the chemokine and relaxin signalling pathways. In Cluster 2, autophagy-related biological processes (Figure 6D and Figure supplement 6C) and pathways were enriched; thus, we defined Cluster 2 as the “autophagy-associated cluster” (AAC) because of its scavenging functions. Finally, Cluster 3 was defined as the “catabolism-associated cluster” (CAC) because of the enrichment of catabolic processes (Figure 6E and Figure supplement 6D) and the Rap1 signalling and RNA degradation pathways. The above results show that m1A modification plays different regulatory roles in the different clusters, indicating the existence of specific m1A modification patterns in neurons. The significance of the modification patterns needs further study.

**Figure 6.**
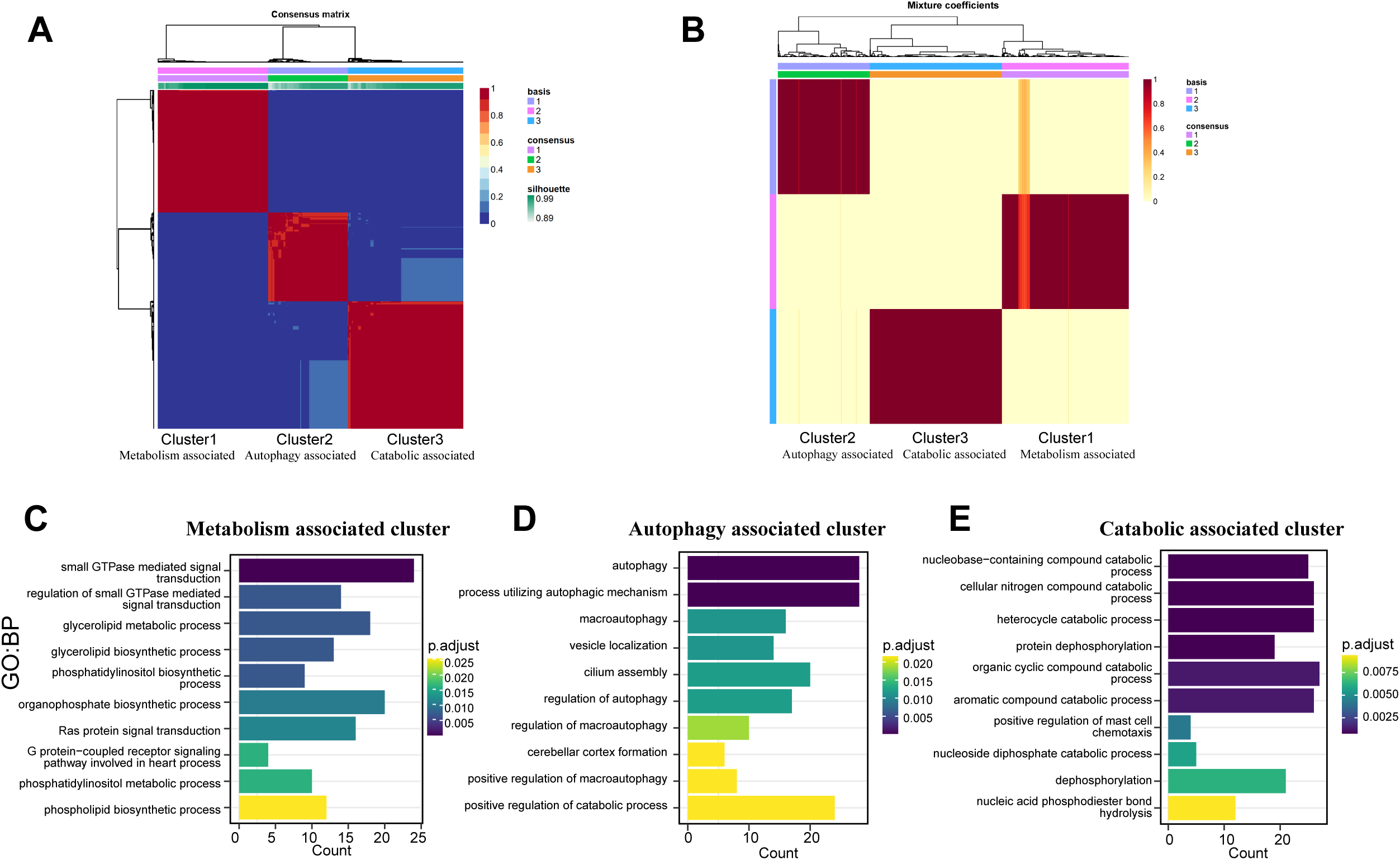
Three patterns regulate m1A modification in neurons. A. NMF analysis shows three m1A modification clusters (termed the metabolism-associated cluster (MAC), autophagy-associated cluster (AAC), and catabolic-associated cluster (CAC)). B. Mixture coefficients matrix also shows that the three clusters have good discrimination. D-E. GO enrichment analysis shows the different biological functions of each cluster. **Figure supplement 6. Three patterns regulate m1A modification in neurons**

## DISCUSSION

m1A is an abundant modification across eukaryotes. Although some studies have reported the landscape and characteristics of m1A modification in tissues and cells *(**Dominissini et al., 2016; Roundtree et al., 2017; Safra et al., 2017**)*, its features and functions in neurons remain unclear. In this study, we identified m1A modifications on three kinds of RNAs (mRNAs, lncRNAs, and circRNAs) in normal neurons and OGD/R-treated neurons. We identified thousands of m1A mRNAs and found that the number and level of mRNA m1A modifications were increased after OGD/R. Regarding lncRNAs, in addition to exploring the basic features of m1A modification, we found two important m1A lncRNAs, Meg3 and Neat1, to be associated with RBPs and to possibly play vital roles in the regulation of mRNA metabolism. There are approximately 3000 m1A circRNAs in neurons, and they play roles in the nervous system. The m1A modifications on circRNAs such as APP and FOXO5 may be involved in the mechanism of circRNA translation. Furthermore, we explored the underlying associations among m1A RNAs (circRNAs, lncRNAs and mRNAs) and found that there are two different complex regulatory networks for differential expression and differential methylation. The m1A modification patterns in neurons were also investigated. Three modification patterns (i.e., clusters) were identified: the metabolism-associated cluster, autophagy-associated cluster, and catabolism-associated cluster. Finally, we confirmed that m1A modification of some mRNAs affects the function of other regulatory mechanisms.

Relatively conserved sequences in RNAs are usually identified as sites of various chemical modifications. Methyltransferases can modify RNAs by recognizing those motifs. There are different motif sequences for different RNA modifications. m6A modification, the most important and common modification from yeast to humans, is located predominantly at RRACU (where R = A/G) consensus motifs in mammals and RGAC (where R = A/G) consensus motifs across yeast species *(**Roundtree et al., 2017**)*. Several studies have identified motifs for m1A modification. The GUUCRA motif was identified in the mitochondrial transcriptome *(**Li et al., 2017**)*. Another study also indicated that m1A was preferentially located at a GA-rich motif in 10-15% of cases *(**Dominissini et al., 2016**)*. The motifs identified in these different studies were inconsistent with our findings. We identified different motifs in neurons, mRNAs (CMGCWGC and GCGGCGGCGGC), lncRNAs (AGARRAARAARAARA and TGCTGCTGCTG) and circRNAs (CWTCNTC and TGGARRA). Some modifications *(**Dominissini et al., 2016; Safra et al., 2017**)*, such as m1A, are more likely to be present in regions with a high GC content, as found in previous studies. In addition, our results show that m1A-modified motifs may have RNA species specificity. These m1A motifs differ greatly across RNAs. Combining our findings with those of a previous study indicating that motifs have RNA methyltransferase specificity *(**Zhang and Jia, 2018**)*, we speculate that different methyltransferases may regulate the methylation of different RNAs. Currently, the results of methylation motif identification seem relatively complex. Further identification of different methylases for different RNAs combined with experimental verification may be a future research direction.

Different RNAs play different roles in biological processes. mRNAs are translated into proteins that regulate biological activities, while noncoding RNAs, such as lncRNAs and circRNAs, play important roles in posttranscriptional regulation via various mechanisms. Chemical modification of these RNAs changes their original regulatory mechanisms and influences their downstream effect. m6A modification of mRNAs can affect mRNA expression, stability, and splicing and can promote or prevent mRNA degradation *(**Tang et al., 2018; Song et al., 2020**)*. m6A modification of circRNAs can affect the translation of circRNAs and splicing of longer circRNAs *(**Wesselhoeft et al., 2019; Huang et al., 2021**)*. Similarly, m6A modification of lncRNAs also plays vital roles in lncRNA-mediated gene silencing and RNA stability *(**Zheng et al., 2019; Yang et al., 2020b**)*. Currently, few studies have addressed the effect of m1A modification on RNA metabolism and functions. Some studies have shown that m1A modification of mRNA can regulate mRNA translation. In addition, we found that m1A modification can affect the ceRNA mechanism by affecting the binding of miRNAs to mRNAs. However, little is known about the function of m1A modification on noncoding RNAs. In this study, we identified the features of m1A modification on lncRNAs and circRNAs and summarized the numbers of genomic sources and intergroup differences in m1A RNAs. Regarding m1A lncRNAs, we explored the RBPs that may interact with m1A lncRNAs and found that the RBPs bound to m1A lncRNAs are involved mainly in the metabolism and splicing of mRNAs. Regarding m1A circRNAs, in addition to drawing a basic map, we also studied the translation characteristics of m1A circRNAs. We found that m1A circRNAs can be translated into proteins and that some of these m1A circRNAs also have m6A-modified sites. The combined findings of current studies *(**Huang et al., 2021**)* indicate that m6A modification regulates circRNA translation. Therefore, if a circRNA simultaneously contains two modifications, which modification has a greater impact on translation regulation, and what is the underlying mechanism? These questions deserve in-depth study.

Transcriptome-wide m1A modification may play a role in some currently identified posttranscriptional regulatory mechanisms. The regulatory mechanism of ceRNAs is generally considered to play a role at the expression level, but the effect of methylation modifications on this mechanism is not clear. In this study, two interaction networks were constructed by using differential expression and differential methylation data, and both networks were extremely complex. Similar regulatory loops between lncRNAs and circRNAs were formed by linking miRNAs, especially in the differential modification regulatory network. Modifications of these RNAs may affect the interactions in these loops. For example, modification of lncRNAs, circRNAs and mRNAs may affect their interactions with miRNAs. This mechanism provides a more accurate strategy for the upstream and downstream regulation of miRNAs, which in turn more conveniently regulates biological processes. In this study, we also found multiple miRNA‒mRNA pairs, and the dual luciferase assay confirmed that m1A modification of some mRNA 3’UTRs affects the binding of miRNAs to these mRNAs. We found that these genes affected by m1A modification (Rbfox3 and Arid1a) are related to neural development and differentiation; the genes that were not affected (Grin2d are Wdtc1) are related mostly to broader cellular functions. Therefore, we asked whether m1A modification preferentially occurs on certain genes. m1A modification depends on several methyltransferases. Thus, do methyltransferases exhibit preferences for different functional genes? Just as we proposed that m1A modification of different RNA species may be mediated by different m1A-related enzymes, we hypothesize that different methyltransferases may exhibit preferences for different functional genes.

In addition, in the exploration of m1A methylation patterns in neurons, as our results showed, m1A modification of different RNAs may be regulated by different enzymes, and different functional genes may be regulated via different methyltransferases; thus, identification of these enzymes is urgently needed. We also identified three patterns of m1A modification in neurons: a metabolism-associated cluster, an autophagy-associated cluster, and a catabolism-associated cluster. The genes with each pattern of m1A modification perform different biological functions. Based on previous studies in other cells *(**Dominissini et al., 2016; Safra et al., 2017; Zhao et al., 2019**)*, we asked whether these m1A modifications are cell type specific. Investigation of this possibility also requires more sequencing and experimental data for analysis.

Overall, in this study, we profiled the features and patterns of m1A modification in neurons and OGD/R-treated neurons. We identified m1A modifications on different RNAs and explored the possible effects of m1A modification on the functions of different RNAs and the overall posttranscriptional regulation mechanism. Although we found and proposed some roles related to m1A modification, we also found that RNA m1Amodification is a relatively complex modification type (no completely conserved modification motif has been found, and m1A may exhibit RNA type specificity, cell type specificity, etc.). Therefore, in-depth study of the importance of this modification for various biological processes and the development of related therapeutic targets based on this modification require considerably more integrated bioinformatic and basic experimental research.

## MATERIALS AND METHODS

### Animals

C57BL6 mice purchased from the Laboratory Animal Centre, Academy of Military Medical Science (Beijing, China), were used mainly for harvesting primary cerebral neurons. All experiments were performed in adherence to the National Institutes of Health Guidelines for the Care and Use of Laboratory Animals and were approved by the Medical Ethics Committee of Qilu Hospital of Shandong University.

### Primary cerebral neuron isolation and culture

Primary cultures of cerebral neurons were obtained as described previously *(**Hilgenberg and Smith, 2007**)*. In brief, mouse foetuses (embryonic day 17 [E17]) were removed from the uterus, and the individual foetuses were freed from the embryonic sacs. The brain and cortical tissue were dissected and placed in high-glucose Dulbecco’s modified Eagle’s medium (DMEM-HG) without phenol red (Gibco, Grand Island, NY, USA; Cat. No. 31053028). Papain solution (10 U/mL; Sigma‒Aldrich, St. Louis, MO, USA; Cat. No. LS003126) was added to these cerebral tissues, which were then incubated for 15 min at 37°C in a 5% CO_2_ incubator. Dissociated cortical cells were plated on poly-L-lysine (Sigma‒Aldrich; Cat. No. P4832)-coated cell culture dishes and cultured in DMEM-HG (Gibco, Grand Island, NY, USA; Cat. No. 31053028) containing 10% foetal bovine serum (Gibco, Australia; Cat. No. 10099141) and 1% penicillin/streptomycin (P/S; Invitrogen, Carlsbad, CA, USA; Cat. No. 15140148) at a density of 1.0 × 10^6^ cells/mL. Four hours after seeding, the medium was replaced with neurobasal medium (NM; Gibco, Carlsbad; Cat. No. 21103049) supplemented with B-27 (Gibco, Grand Island, NY, USA; Cat. No. 17504044). Cells were cultured in a humidified incubator at 37°C with 5% CO_2_. The medium was changed every 3 days. Cultures were used for in vitro experiments after 7 days.

### OGD/R modelling

The OGD/R model was established using a previously described method with slight modifications *(**Tasca et al., 2015; Ryou and Mallet, 2018**)*. Cultured primary cerebral neurons were washed twice with phosphate-buffered saline (PBS; Sigma‒Aldrich; Cat. No. D8537) supplemented with 1% P/S after 7 days of culture. Glucose-free Dulbecco’s modified Eagle’s medium (Gibco, Grand Island, NY, USA; Cat. No. 31053028) was added to the dishes. Next, the neurons were cultured with a GENbag anaerobic incubation system (bioMérieux SA, France; Cat. No. 45534) at 37°C. The cultures were kept separate under hypoxic conditions for 1.5 and 3 h to achieve OGD. Thereafter, the neurons were allowed to recover by culture in normal serum-free medium (NM) under normal incubation conditions (37°C, 5% CO_2_) for 24 h. Neurons cultured in normal serum-free medium under normoxic conditions served as controls.

### RNA isolation

Total RNA was extracted from primary cultured neurons using TRIzol reagent (Invitrogen, Carlsbad, CA, USA; Cat. No. 15596018) according to the manufacturer’s protocol. A NanoDrop ND-1000 system (Thermo Fisher Scientific, Waltham, MA, USA) was used to measure the RNA concentration in each sample. The OD260/OD280 ratio was assessed as an index of RNA purity, and samples with OD260/OD280 values ranging from 1.8 to 2.1 met the qualifications for purity. RNA integrity was evaluated using denaturing agarose gel electrophoresis.

### Preparation of the m1A RNA immunoprecipitation sequencing (MeRIP-seq) library

MeRIP-seq was performed by Cloudseq Biotech Inc. (Shanghai, China) according to a published procedure with slight modifications *(**Meyer et al., 2012**)*. In brief, three biological replicates were used for the control, OGD/R 1.5 h, and OGD/R 3 h groups. Isolated RNA was chemically degraded into fragments of approximately 100 nucleotides in length using fragmentation buffer (Illumina, Inc., CA, USA). A Magna MeRIP™ m1A Kit (Merck Millipore, MA, USA; Cat. No. 17–10499) was used to perform immunoprecipitation (IP) of m1A RNA according to the manufacturer’s recommendations. IP buffer (Cat. No. CS220009), Magna ChIP Protein A/G Magnetic Beads (Cat. No. CS203152) and an anti-m1A antibody (Cat. No. MABE1006) were used to prepare the magnetic beads for IP. Next, the MeRIP reaction mixture was prepared according to the manufacturer’s guidelines, and fragmented RNA was included in this mixture. The magnetic beads and MeRIP reaction mixture were combined in tubes, and all tubes were incubated with rotation for 2 h at 4°C. Subsequently, elution buffer was prepared according to the manufacturer’s instructions and was used to elute bound RNA from the beads using the anti-m1A antibody in IP buffer. Both input samples without IP and m1A input samples were used for library construction using an NEBNext® Ultra™ II Directional RNA Library Prep Kit (New England Biolabs, Inc., MA, USA). Eluted RNA fragments were converted to cDNA and sequenced or treated to induce partial m1A to m6A conversion before cDNA synthesis. Library sequencing was performed using an Illumina HiSeq 4000 instrument (Illumina, Inc., CA, USA) with 150-bp paired-end reads.

### MeRIP-seq data analysis

In brief, quality control of the paired-end reads was performed with FastQC (v0.11.9) prior to trimming of 3’ adaptors and removal of low-quality reads using Cutadapt software (v1.9.3). Then, HISAT2 software (v2.0.4) was used to align the clean reads from all libraries to the reference genome (mm10) downloaded from Ensembl. Peaks for which -10×log10(*p* value) > 3 were detected using Model-Based Analysis of ChIP-Seq (MACS) software (v2.2.7.1). Differentially methylated sites with a fold change of ≥ 2 and false discovery rate (FDR) of ≤ 0.0001 were identified with the diffReps differential analysis package (v1.55.6). The peaks identified by MACS and diffReps that overlapped with exons of mRNAs, lncRNAs, and circRNAs were identified and selected for further analysis. In addition to conventional peak calling (conventional method), the peaks with A->T mismatches (mismatch method) were analysed. In addition, m1A sites have a partial chance of terminating reverse transcription, resulting in lower coverage in the middle of the peak (near the m1A site) than on both sides, forming a depression. These peaks and depressions (trough method) were also analysed. Moreover, when m1A was converted into m6A, the mismatch and termination properties of m6A were lost, while reverse transcription was normal. Thus, after sequencing, peaks with the normal shape could be detected at the sites (treatment method).

### Preparation of RNA sequencing (RNA-seq) libraries

Total RNA was used for removal of ribosomal RNA (rRNA) using Ribo-Zero rRNA Removal Kits (Illumina, USA) following the manufacturer’s instructions. RNA libraries were constructed by using rRNA-depleted RNAs with a TruSeq Stranded Total RNA Library Prep Kit (Illumina, USA) according to the manufacturer’s instructions. Libraries were subjected to quality control and quantified using a BioAnalyzer 2100 system (Agilent Technologies, USA). RNA was purified from each sample using Oligo(dT) Dynabeads (Invitrogen) and subjected to first-strand cDNA synthesis and library preparation using a TruSeq Stranded mRNA Library Prep Kit (Illumina). Libraries (10 pM) were denatured as single-stranded DNA molecules, captured on Illumina flow cells, amplified *in situ* as clusters and finally sequenced for 150 cycles on an Illumina HiSeq 4000 instrument according to the manufacturer’s instructions.

### RNA-seq data analysis

Quality control of paired-end reads was performed with FastQC (v0.11.9) prior to trimming of 3’ adaptors and removal of low-quality reads using Cutadapt software (v1.9.3). The high-quality reads were aligned to the mouse reference genome (UCSC mm10) with HISAT2 software (v2.0.4). Then, guided by the Ensembl (GRCm39.104) GTF gene annotation file, expression was estimated in units of fragments per kilobase of transcript per million mapped reads (FPKMs). The differentially expressed genes were identified as those with a fold change of ≥ 2 and adjusted *p* value of ≤ 0.05.

### Preparation of miRNA sequencing (miRNA-seq) libraries and data analysis

miRNA-seq was conducted at Cloudseq Biotech Inc. (Shanghai, China). In brief, total RNA from each group was prepared and quantified with the BioAnalyzer 2100 system (Agilent Technologies, USA). Polyacrylamide gel electrophoresis (PAGE) was performed, and the gel was cut to select the band corresponding to a length of 18-30 nt to recover small RNAs. Adaptors were then ligated to the 5’ and 3’ ends of the small RNAs. After cDNA synthesis and amplification, the PCR-amplified fragments were purified from the PAGE gel, and the complete cDNA libraries were quantified with a BioAnalyzer 2100. Sequencing was performed on the Illumina HiSeq 4000 instrument, and 50 bp single-end reads were generated. For miRNA-seq data analysis, the adaptor sequences were trimmed, and the trimmed reads (≥ 15 nt) were retained by Cutadapt software (v1.9.3). Then, the trimmed reads were aligned to the merged mouse pre-miRNA databases (known pre-miRNAs from miRbase [v22.1]) using NovoAlign software (v3.02.12). The number of mature miRNA-mapped tags was defined as the raw expression level of that miRNA. Read counts were normalized by tag counts per million aligned miRNAs (TPM) values. Differentially expressed miRNAs between the two groups were filtered by the following criteria: fold change ≥ 2 and adjusted *p* value ≤ 0.05.

### Competing endogenous RNA (ceRNA) regulatory network construction

Based on the miRNA expression data, we used three databases (TargetScan, miRDB, and miRTarBase) for comprehensive analysis of miRNA binding target genes. We obtained the intersection of the mRNAs predicted by the three databases, our differentially expressed mRNAs and our differentially methylated mRNAs to construct the miRNA‒mRNA interaction network. For lncRNAs and circRNAs, we used the ENCORI database (https://starbase.sysu.edu.cn/) to analyse the binding of lncRNAs and circRNAs to the differential miRNAs. After obtaining the lncRNA‒miRNA and circRNA–miRNA interaction networks, we intersected the included RNAs with the differentially expressed lncRNAs and circRNAs and the differentially methylated lncRNAs and circRNAs in our data. Finally, we integrated all data to construct the expression and methylated circRNA/lncRNA‒miRNA‒mRNA interaction networks.

### Nonnegative matrix factorization (NMF) clustering

NMF is a widely used tool for the analysis of high-dimensional data because it automatically extracts sparse and meaningful features from a set of nonnegative data vectors. NMF clustering was used to determine the m1A modification patterns in our 9 differently treated samples *(**Gaujoux and Seoighe, 2010**)*. We performed this analysis using all m1A-modified mRNAs in the three groups. The *k* value at which the magnitude of the cophenetic correlation coefficient began to decrease was selected as the optimal number of clusters. The heatmap of m1A regulators, the basis components, and the NMF connectivity matrix for different clusters were estimated with the NMF package (v0.24.0) in R (v4.1.3).

### Functional annotation

Gene Ontology (GO) and Kyoto Encyclopedia of Genes and Genomes (KEGG) pathway enrichment analyses based on the differentially expressed genes were performed with the R package clusterProfiler (v4.4.0) *(**Wu et al., 2021**)*. GO covers three categories: cellular component (CC), molecular function (MF), and biological process (BP). The *p* value for a GO term denotes the significance of enrichment of genes in that term. Pathway enrichment analysis is a functional analysis that maps genes to KEGG pathways. The Fisher *p* value denotes the significance of the pathway correlation to the conditions. The GO and KEGG pathway terms with adjusted *p* values of ≤ 0.05 were considered significantly enriched.

### Luciferase activity assay

The Rbfox3, Arid1a, Grin2d and Wdtc1 reporter vectors were constructed by inserting the 3’UTRs of the corresponding genes downstream of the luciferase reporter gene into the GV272 plasmid (GeneChem Technologies, Shanghai, China). The Trmt6 overexpression vector was constructed by subcloning the coding sequence (CDS) into the GV712 plasmid (GeneChem Technologies, Shanghai, China). The miR-129-5p, miR-101b-3p, miR-27-3p and miR-16-5p mimics were designed and synthesized by GeneChem. We used Lipofectamine 3000 to cotransfect reporter plasmids, the Trmt6 plasmid and miRNA mimics into HEK293T cells. Luciferase activity was measured 48 h later according to the manufacturer’s procedures (E292, Promega, USA). All experiments described were replicated independently with similar results at least three times.

## AUTHOR CONTRIBUTIONS

Hengxing Zhou, Shiqing Feng, Mingjun Bi and Xianfu Yi designed the experiments and edited the paper; Chi Zhang performed the experiments, analysed the data and drafted the manuscript and figures; Xianfu Yi and Mengfan Hou analysed the data and drafted figures; Qingyang Li and Xueying Li participated in the design of the study and contributed to the isolation of RNA; Lu Lu, Enling Qi and Mingxin Wu contributed to the OGD/R model; Lin Qi and Huan Jian contributed to the sequencing data analysis; Zhangyang Qi contributed to the MeRIP qRT-PCR experiment; and Yigang Lv and Xiaohong Kong helped to design the experiment and interpret the data. All authors read and approved the final manuscript.

## ACKNOWLEDGEMENTS

We also thank Dr Jianming Zeng (University of Macau), and all the members of his bioinformatics team, biotrainee, for generously sharing their experience and codes.

## DATA AND CODE AVAILABILITY

The datasets generated and analysed in the current study are available in the BIGD (National Genomics Data Center, Beijing Institute of Genomics, Chinese Academy of Sciences) database under the bioproject accession code PRJCA010978. No custom code was generated, name and version of the softwares are included in the method.

